# Uncovering Pseudotime-Varying Genetic Causal Effects Along Single-Cell Trajectories for Pulmonary Disease Trait

**DOI:** 10.64898/2026.06.08.730759

**Authors:** Siyuan Chen, Ahila Moorthy, Paul Kwong Yu, Jingshu Wang, Dajiang Liu

## Abstract

With the increasing accessibility of single-cell RNA sequencing (scRNA-seq) data, cell-type-specific gene expression can be linked to complex traits through pseudo-bulk method, which considered aggregated gene expression from multiple cells of the same annotated cell type per individual and clearly shows the limitation of ignoring intra-individual cell-to-cell variability. Concurrently, pseudotime trajectory inference has gained popularity for its ability to capture continuous biological processes such as cell differentiation and lineage development, instead of individual discrete stages. It is natural to consider whether genetic effects for complex traits, such as individual level disease status, show a dynamic pattern along the inferred trajectories. In this study, we introduce a novel framework that models gene expression as a function of pseudotime along the inferred trajectories. We mapped expression quantitative trait loci (eQTL) effects in the cis-region as functional parameters, which we called “dynamic eQTLs”, showing regulatory effects exerted by genetic variants change continuously along the cellular trajectory. For eQTLs of constant effects across pseudotime we leveraged external bulk-eQTL information to enhance the power. Furthermore, we employed significant, variable dynamic eQTLs as instrumental variables to infer causal relationships between gene expression and complex traits. To address challenges inherent to scRNA-seq data—such as sparsity and high variability—we incorporate an empirical likelihood-based inference method, which is non-parametric and self-normalized. Besides, genes associated with trajectory branchpoints may bring confounding, and we also proposed a causal mediation analysis framework to determine whether a gene plays a causal role for the disease directly and indirectly through driving cell fates. Applying our method to scRNA-seq data from human lung tissue of 114 samples (66 pulmonary fibrosis cases and 48 controls), along with meta-analyzed GWAS summary statistics for IPF from 3 studies, we identified pseudotime-dependent causal effects for IPF from genes implicated in the trajectory AT2 – translational AT2 – AT1, which is crucial in lung tissue repair and regeneration. We also found that 30 genes have a mediated effect through cell fates.

## Introduction

Single-cell RNA sequencing (scRNA-seq) has enabled the high-throughput quantification of gene expression specific to cell types and states. With the cost of scRNA-seq decreasing and techniques for sample multiplexing improving, population-scale scRNA-seq, and thus single-cell expression quantitative trait locus (sc-eQTL) mapping, is increasingly feasible. Statistical methods for bulk eQTL mapping, such as linear mixed effect models, have been implemented for annotated homogeneous cell populations, i.e., the “pseudo-bulk” method. The major advantages of this approach are that cell aggregation can mitigate sparsity, while the following limitations exist [1]. Firstly, it is challenging to determine the appropriate resolution to model discrete cell states without losing signals of that some genes differentially expressed. Besides, upon aggregation, intra-individual cell-to-cell variability of the same cell type label is ignored. To overcome these problems, single-cell models that treat the expression for each cell as an observation offer a strategy for unbiased identification of cell-state-specific regulatory effects. This is especially relevant when dynamic effects may be present in more granular cell states or along a continuous cell-state transition. Obviously, statistical methods for (pseudo-)bulk models are not applicable anymore here. Raw gene expression counts or log-transformed expression in each cell for a gene can be modelled including donor-level and cell-level fixed effects, and random effects from repeated measurement from the same donor [2].

When cells in a sample come from a continuous biological process, computationally placing the cells along a pseudo-temporal trajectory based on their progressively changing transcriptomes is a powerful approach to reconstructing the dynamic gene expression programs of the underlying biological process. This approach, also known as pseudotime analysis, is now widely used to study cell differentiation, immune responses, disease development, and many other biological systems with temporal dynamics. While trajectory inference, especially the methods based on minimum spanning trees such as Slingshot [3], aims for establishing a branching structure based on a dimension-reduced space of cell by gene expression matrix, the pseudotime is a distance metric from some origin for cells, rather than the actual concept of time [4]. Computationally inferred trajectories can be validated by known cell differentiation procedures. Naturally, pseudotime defined on a known trajectory serves a good metric for the concept of “continuous cell states”, as it clearly has a biological meaning that measures similarity among cells and ambiguity in cell type annotation for each cell. While treating cell states as a continuous variable rather than discrete categories is not a new idea. In general, the benefits of considering cell types as continuous are the following. Firstly, it reduces the inference bias brought by the ambiguous boundary cells, and this becomes more important for relatively rare cell types. Besides, it enables a dynamic perspective for effects like eQTL. For example, OneK1K [5] reports eQTL effect change from the 1^st^ to the 3^rd^ quantiles of the trajectory from naïve B cells to memory B cells. This reminds us that genetic effects can vary within the same cell type rather than just measurement errors. Finally, it is a new option to account for cell-to-cell variability, which could be better than just considering it as a random effect, since such variability could be too fierce to be assumed following a Normal distribution of zero mean and a fixed variance.

Aside from eQTL mapping from the perspective of continuous cell types, we are motivated to establish a complete causal inference framework for causal gene discovery based on our observation when working on the scRNA-Seq data from human lung from [6]. The association between gene expression and pulmonary diseases, such as lung cancer, chronic obstructive pulmonary disease (COPD), and various forms of pulmonary fibrosis, has been explored using single-cell RNA-Seq data through methods such as differential expression analysis. A molecular atlas of the human lung identified 58 distinct cell types, including 41 previously known and 14 newly discovered. Changes in cell type populations and gene enrichment have been observed in disease samples. However, a clear causal relationship between gene expression in specific cell types and disease status has yet to be established. This lack of clarity arises due to several challenges: (1) cell type identification typically relies on known marker genes, but over- or under-expression of certain genes may drive cell type transitions, and (2) gene expression within closely related cell types is often highly correlated, making it difficult to isolate individual gene effects; (3) there could be topological differences, such as a cell lineage along differentiation is lost (or added) in the disease samples comparing to the controls [7].

Our goal is to develop a comprehensive causal model to elucidate the role of cell type-specific gene expression, particularly quantitative trait loci, in lung diseases, along with including the possible effects brought by cell transition “decisions”. This approach will enhance our understanding of lung disease mechanisms and potentially identify novel drug and therapeutic targets. Applying our method to scRNA-seq data from human lung tissue of 114 samples (66 pulmonary fibrosis cases and 48 controls), along with meta-analyzed GWAS summary statistics for Idiopathic Pulmonary Fibrosis (IPF) from 3 studies, we identified pseudotime-dependent causal effects for IPF from 8 genes implicated in the trajectory AT2 – translational AT2 – AT1, which is crucial in lung tissue repair and regeneration. We also found that some of the causal genes have a mediated effect through cell fates at the trajectory branchpoint. Together, these findings highlight the utility of integrating pseudotemporal genetic regulation with cell-state transitions to provide a causal framework for dissecting the molecular mechanisms of IPF.

## Results

### Method Overview

We proposed a framework to integrate the single cell eQTL model and pseudotime trajectory inference based on the idea that cell states or types are continuous and hence the eQTL effect could be varying by the continuous cell states as well. To be more specific, given an inferred trajectory and ordered cells along the trajectory with their pseudotime, gene expression measured in each cell is a function of the pseudotime, while the genotype for each donor remain unchanged along the pseudotime axis. The single-cell model also includes individual-level covariates, such as sex, age, and ancestry PCs for donors, and cell-specific covariates, such as expression-based PCs, as fixed effects, and an individual level random effect term [2]. Most importantly, fitting this model is equivalent to doing a functional-on-scalar regression, which is well-established in the field of functional data analysis. The gene expression by pseudotime observations are firstly smoothed with B-splines of rank decided by the number cell types on the trajectory. From our observation in practice, we recommend using a rank around K(K-1), K is the number of cell types. Then we fit the model to obtain the estimation for the functional parameter for genotype as our pseudotemporal eQTL. The significance of pseudotemporal eQTL is from testing the null hypothesis: *β*(*t*) = 0. Moreover, we mapped pseodotemporal eQTL that has an interaction with disease status of donors by including a functional coefficient for the interaction term.

Based on the mapped pseudotemporal eQTLs, we constructed a Mendelian Randomization framework for detecting causal genes for the individual-level disease status outcomes, as well as whether this causal effect is mediated by cell fates. The eQTLs selected as instrumental variables (IVs) should fulfill the usual IV assumptions [8]: 1. Relevance, this could be achieved by filtering for significant eQTLs; 2. Exchangeability, this is not testable; 3. Exclusion restrictions, violation for this is commonly referred as horizontal pleiotropy, and can be managed through modelling. Besides, the eQTLs with significant interaction with disease status were ruled out as they violated the IV assumption of homogeneous effects. To detect pseudotime-varying causal effects of gene expression, it is necessary that at least one instrumental variable exhibits variation across the pseudotime trajectory. As no formal statistical test currently exists for this criterion, we address it pragmatically by visually inspecting the eQTL trajectories and regulating their smoothness.

Under a two-sample Mendelian Randomization setting, we denote the estimated association between a genetic instrument and the outcome as 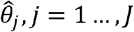. The functional causal effect of a gene on the disease outcome is estimated through the following weighted regression of inverse-variance-weighting method, with an Egger-type horizontal pleiotropy term:

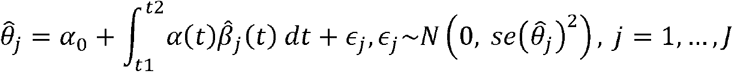

This regression can be fitted using a weighted empirical method [9] (details in Materials and Methods).

Besides, some genes in a cell may express differently at some stage on the trajectory, especially for cells near or at the observed trajectory branchpoint, which may affect cell fates in the way that the cell is more likely to differentiate into one type rather than the other. Fates of multiple cells triggered by gene expression may have a causal effect on the disease outcome trait, which can be regarded as a mediated effect from the gene expression. To explore potential mediation effects, we first choose a subset of cells that appear to be in a small window around the trajectory branchpoint and model the gene expression at the timepoint with a summary-data based MR approach. Such inference can be made pointwise, based on genetic instrument that found by running an association analysis between genetic variants and “cell fate decisions”. Note that since we only consider a very small window for pseudotime in this analysis, a functional exposure is not feasible in this scenario. Hence, we are only testing the sharp null hypothesis *H*_*0*_ : *τ* (*t*_0_ ) = 0 where *t*_0_ denotes the pseudotime value for the branchpoint, *τ* (*t*_0_ ) denote the direct effect of the gene on the disease outcome.

The overall analytical workflow proceeded as follows: (1) identification of cell types and trajectories of interest; (2) dimensionality reduction of the gene expression matrix for cells belonging to the selected types, followed by trajectory inference to obtain pseudotime (e.g., using Slingshot in this study); (3) mapping of pseudotime-dependent eQTLs along the inferred trajectories; and (4) causal inference leveraging external GWAS summary statistics for the outcome of interest.

### Functional pseudotemporal eQTLs reflects transcriptomic effects on biological procedures

We use the data from [6] and look at the trajectory of epithelial cell types AT2 – Transitional AT2 – AT1, inferred and discussed in [10] [11], which has great meaning in alveolar regeneration and repair. To be more specific, [12] demonstrated that following alveolar injury, AT2 cells adopt an intermediate transcriptional state before fully converting to mature AT1 cells, a process mediated by TGF-β signaling. The presence of transitional AT2 cells in the human lung was highlighted in [10], characterized by transitional markers indicative of active remodeling and alveolar regeneration. The other trajectory, as a comparison, is Secretory SCGB3A2+ – Translational AT2 – AT1. We first obtain the values for pseudotime *t* for each cell using Slingshot based on 69463 cells limited to AT2, Transitional AT2, AT1 and Secretory SCGB3A2+ cell types, with starting cluster set as AT2 and Secretory SCGB3A2+. We obtained the same 2 trajectories as [11] (Graph S1). Next we map pseudotemporal eQTLs on these two trajectories separately (details in Materials and Methods). We set the significance level p-value threshold as 1x 10^−9^.

We highlight the following findings. Major of the “pseudotemporal eGenes”, that are the genes with significant pseudotemporal eQTLs, are different between the two trajectories. Because the number of available cells for each trajectory differ, the number of eGenes are not directly comparable. We focus on the eGenes shared by both trajectories, and the eGenes on AT2 – Transitional AT2 – AT1 and demonstrate their biological relevance with lung repairment and regeneration. (1) In the intersection of eGenes there are 106 genes. GO enrichment analysis of these genes revealed enrichment of fundamental cellular processes such as ion homeostasis, stress responses, platelet activation, and extracellular matrix organization (Figure 1). These results suggest that the identified genes are broadly relevant to maintaining epithelial cell integrity, response to injury, and remodeling—consistent with central regulatory or ‘hub’ functional roles within lung tissue. (2) GO enrichment analysis of eGenes (Figure 1) along the AT2–transitional AT2–AT1 trajectory indicates that, in addition to their role in epithelial differentiation, these genes are also implicated in immune regulatory processes, including macrophage proliferation, chemotaxis, and dendritic cell antigen processing, consistent with the recognized function of AT2 cells in communication with resident immune cells during homeostasis and injury [13]. Additionally, enriched processes related to retinoic acid and elastic fiber assembly highlight active tissue remodeling and potential involvement in fibrotic pathways [14]. Moreover, Enrichment of stress responses to metal ions likely reflects adaptive mechanisms mitigating oxidative stress during epithelial differentiation, while enrichment of negative regulation of SMAD phosphorylation points to intrinsic control of TGF-β–driven profibrotic signaling. Collectively, these genes are indicative of essential regulators orchestrating immune response, repair, and homeostatic mechanisms within lung epithelial cell differentiation. (3) Of the 271 eGenes on AT2–Transitional AT2–AT1, 52 carry variants that markedly disrupt binding motifs of transcription factors with established roles in lung and epithelial lineage specification (e.g., THRA[15], KLF5, NFIB [16], HEY1[17], ETV5[18]) with a stringent threshold ( *p* 1 x 10^−4^). These genes are integral to a wide array of biological processes essential for pulmonary health and disease. For instance, genes like *SPARC, COL17AA1, SERPINA3*, and the heat shock proteins *HSPA1A/B* are well-established regulators of epithelial integrity and cellular stress responses. Functional variants within these genes can compromise the structural resilience and adaptive capacity of the lung epithelium. The list also includes key mediators of immune signaling, such as *IL6, TNFSF14*, and *SERPING1*, where genetic disruptions can alter inflammatory responses, a cornerstone of many pulmonary pathologies. Furthermore, a significant number of genes are involved in extracellular matrix remodeling (e.g., *ABI3BP, ADAMTS9, DKK1, DKK3, FGF7*), suggesting that variants affecting their regulation by transcription factors could influence the structural reorganization that occurs during alveolar differentiation and repair. This is complemented by genes involved in metal ion binding and detoxification (e.g., *MT1E, MT1M, CPVL, LTF*), which are part of the stress-adaptive programs active in transitional epithelial states. The list is further populated by genes associated with the cell cycle and proliferation (e.g., *CCND2, CKS2, MAPK15*) and a diverse array of signaling and receptor molecules (e.g., *KCNMA1, SLC6A4, RAMP2, PLPP1*), highlighting the multifaceted impact of these regulatory variants on lung cell biology. Together, these results indicate that a diverse set of functional modules—encompassing extracellular matrix regulation, immune modulation, stress and ion homeostasis, and epithelial growth—may be under strong genetic influence through TF binding disruption in the AT2–AT1 differentiation trajectory.

**Figure 1.**
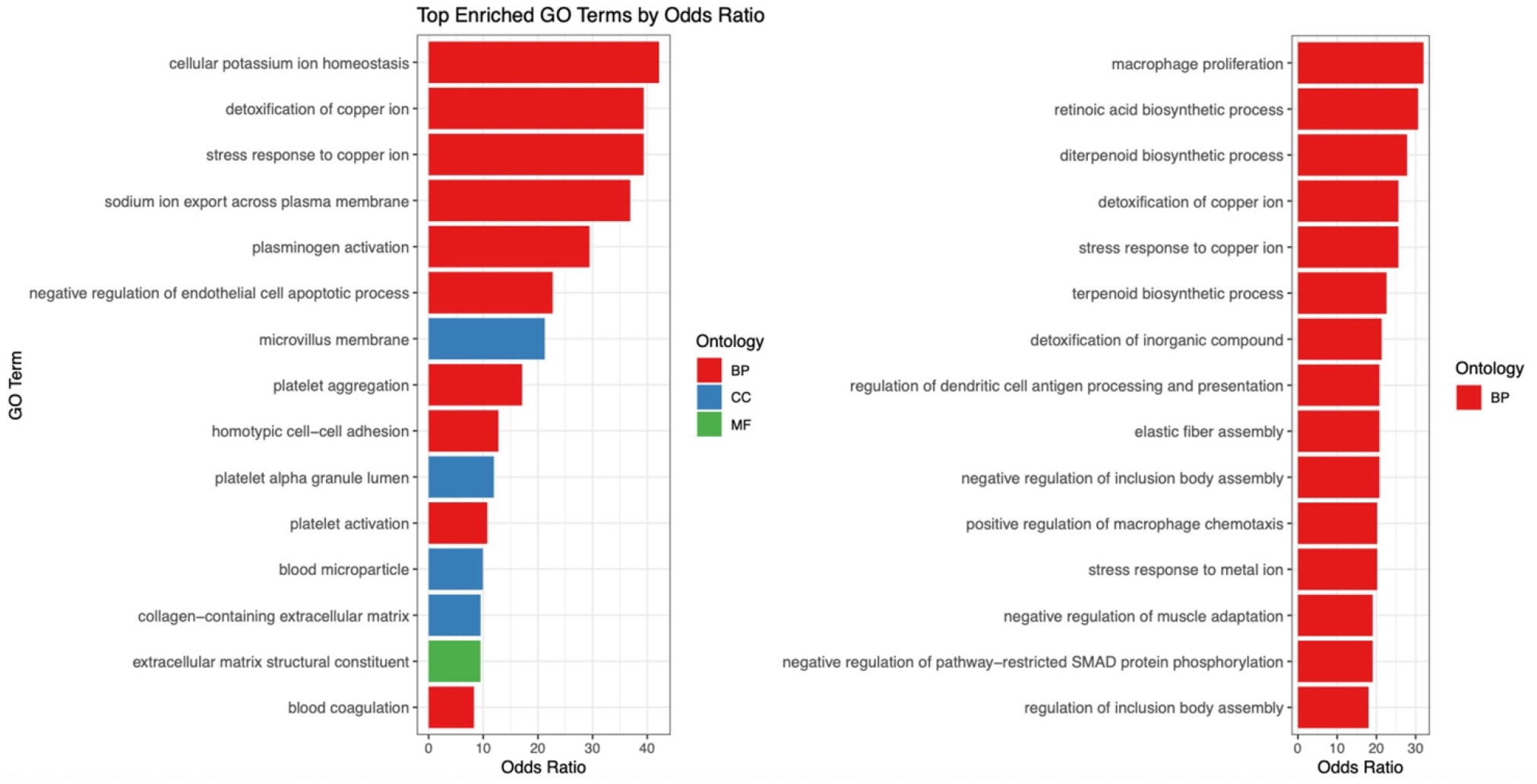
Top GO enriched terms for shared eGenes (left) and AT2-transitional AT2-AT1 trajectory alone (right).

**Figure 2.**
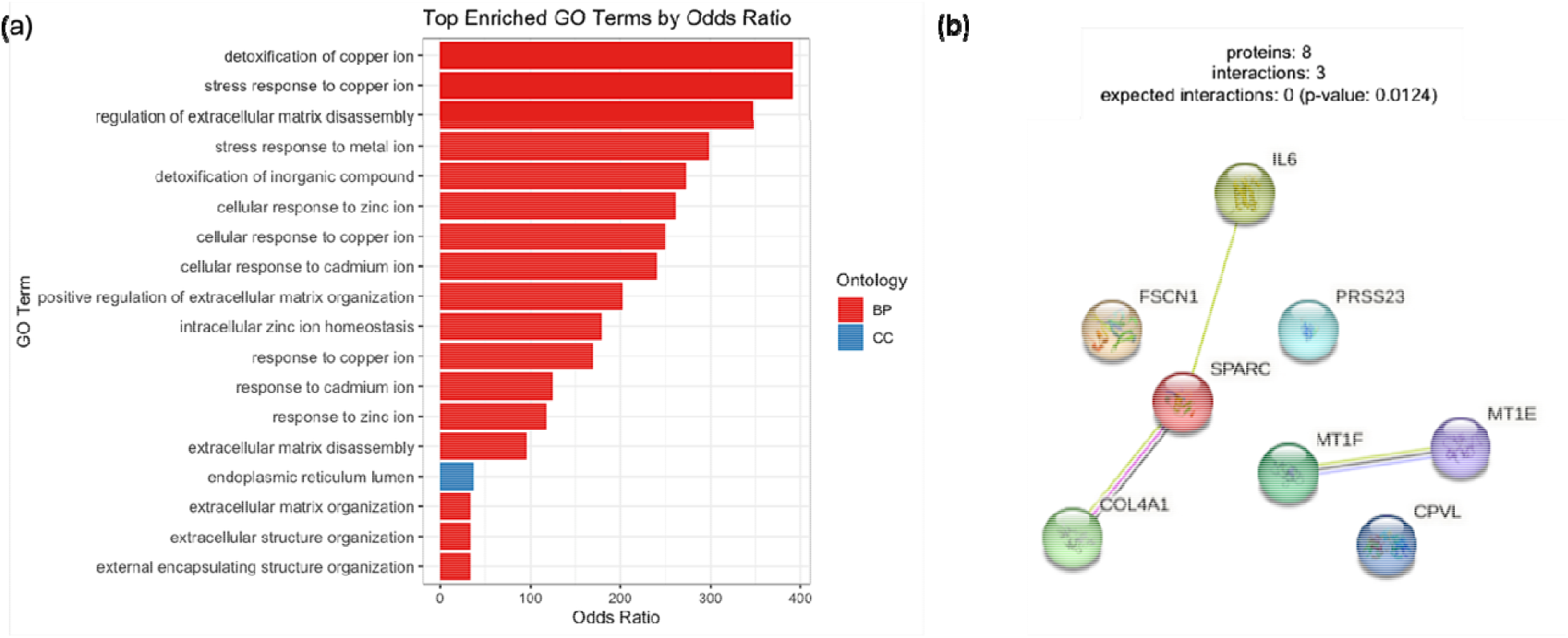
(a) Top GO terms for causal genes on trajectory AT2-transitional AT2-AT1. (b) STRING PPI network for causal genes.

Among the eQTL identified to interact with idiopathic pulmonary fibrosis (IPF) disease status (significance level 1e-5), several genes highlight well-established pathogenic pathways while also nominating novel candidates of potential relevance. Strongly supported immune mediators include *CCL18* and *CD163*, both linked to profibrotic macrophage activation and fibroblast recruitment, and structural regulators such as *CDH11* and *COL11A1*, which drive extracellular matrix remodeling and myofibroblast differentiation. In addition, functionally plausible but less characterized signals were observed at *ALDH1A2*, a retinoic acid–synthesizing enzyme critical for alveolar repair, and *BAMBI*, a negative regulator of TGF-β signaling that may restrain profibrotic activity. Lipid and oxidative stress– related genes such as *ALOX15* and *APOD*, as well as matrix-remodeling factors like *CEMIP* and *ABI3BP*, further underscore the convergence of metabolic, epithelial, and stromal dysregulation in the fibrotic lung.

### Calibration results

We use the bulk eQTL results from TOPMed [19] to calibrate the constant eQTL effects on given trajectory (details in Methods and Materials). Our calibration method managed to identify novel significant constant eQTLs, as shown in Table 1. P-values are BH corrected. We also conducted GO enrichment analysis for genes with significant constant eQTLs for each trajectory. The GO enrichment analysis of genes with constant eQTL effects along the AT2–transitional AT2–AT1 trajectory reveals a strong enrichment for immune-related processes, including macrophage chemotaxis, leukocyte migration, and acute-phase response, showing similar pattern as discussed before. Additionally, terms fundamentally related to epithelial structure and cytoskeletal organization—such as adherens junction organization, axoneme assembly, and microtubule bundle formation—indicate persistent control over structural remodeling required for epithelial cells. Enrichment of apoptosis-regulating pathways and processes like regulation of body fluid levels further highlights the role of these genes in maintaining epithelial homeostasis and barrier function. These results suggest that constant eQTLs preferentially regulate core immune-epithelial programs that remain active across alveolar epithelial state transitions, independent of dynamic transcriptional changes.

**Table 1.** Number of significant eQTLs with constant effect over the pseudotime trajectories after calibration.

| Trajectory | # novel significant eQTLs | # total significant eQTLs | Top genes |
| --- | --- | --- | --- |
| AT2 – Transitional AT2 – AT1 | 37 | 134 | *C1S*, *AIF1*, *DNALI1*, *MEST*, *CAPSL*, *CREB3L1*, *HMOX1*, *IFI27*, *PDLIM3*, *SFN* |
| Secretory SCGB3A2+ – Transitional AT2 – AT1 | 46 | 188 | *AIF1*, *CD52*, *LTF*, *DRC1*, *EMP2*, *HPGD*, *ITGA1*, *CDH11*, *DRC3*, *LMCD1*, *LYZ*, *MEST* |

### eQTL results for the OneK1K cohort

We use the data from [5] and look at the trajectory of naïve B cell – memory B cell, which is also discussed in the section “dynamic eQTL”of [5]. To have better computational efficiency, we reduce the model complexity by change the individual-level random effect to cluster-level and donors are firstly clustered by K-means. Out of 404 the most variably expressed genes we found 116 to have significant pseudotemporal eQTLs on the given trajectory. GO enrichment analysis of these genes reveals a strong enrichment of MHC class II-related functions, including antigen processing, peptide loading, and receptor activity. These findings highlight the transcriptional and genetic regulation of immune presentation machinery during B cell maturation. Enriched terms related to MHC protein complex assembly, binding, and ER chaperone activity suggest that memory B cells gain enhanced competency in antigen processing and presentation. The enrichment of both molecular function and cellular component terms associated with MHC class II underscores the pivotal role of memory B cells in adaptive immune surveillance and secondary immune responses.

### Causal genes for IPF

We use the meta-analyzed GWAS summary statistics for IPF on European population from [20][21][22]. We define a gene to be causal in a rather conservative strategy: call a gene significantly causal only when it has non-zero pointwise 95% PI not covering 0 for at least 10% of the time points on the fixed trajectory. The causal genes on AT2 – Transitional AT2 – AT1 trajectory are *SPARC, PRSS23, MT1E, MT1F, IL6, FSCN1, CPVL, COL4A1*, while there is only 1 gene, *EDNRB* found to be significant on the Secretory SCGB3A2+ – Transitional AT2 – AT1 trajectory. Some of these genes, including *SPARC, IL6, EDNRB*, are already connected with the prevalence of IPF, see [23][24][25]. Especially, SPARC is matricellular protein and is known to modulate collagen assembly & TGF-β activity. *PRSS23* is an activator of endothelial-to-mesenchymal transition (EndMT) and contributes to increased extracellular matrix (ECM) protein production [26]. *FSCN1, CPVL, COL4A1* were reported to influence epithelial-to-mesenchymal transition (EMT) [27][28][29], which is believed to contribute to epithelial dysfunction, failed regeneration, and profibrotic signaling loops in IPF pathogenesis. *MT1E* and *MT1F* belongs to the metallothionein family, that was reported to be protective against lung injuries [30] and play a known role of responding to oxidative stress implicated in lung injury/fibrosis.

To investigate the functional relevance of genes identified as causal for idiopathic pulmonary fibrosis (IPF) along the AT2–transitional AT2–AT1 trajectory, we performed GO enrichment analysis. The top enriched biological processes (BP) prominently reflect metal ion detoxification and stress response pathways, including detoxification of copper ion, stress response to copper ion, and response to cadmium ion, implicating a role for metal homeostasis and cellular stress adaptation in the pathogenic transition of alveolar epithelial cells. Additional enriched terms, such as regulation of ECM disassembly and positive regulation of ECM organization, suggest that active remodeling of the ECM is a key regulatory procedure for lung fibrosis. We also observed enrichment in cellular components (CC) such as platelet alpha granule lumen and basement membrane, further supporting the involvement of matrix remodeling and epithelial–mesenchymal crosstalk. Together, these findings indicate that genetic perturbations contributing to IPF may converge on processes that govern cellular responses to metal-induced oxidative stress and structural reorganization of the alveolar niche, which are critical during epithelial injury and repair.

We also conducted protein-protein interaction analysis. The protein–protein interaction (PPI) network comprises eight proteins with three observed interactions, significantly more than expected by chance (p = 0.0124), indicating functional connectivity among a subset of the genes. The network shows two small interaction clusters. One cluster centers on *SPARC*, which interacts directly with *COL4A1* and *IL6*, linking extracellular matrix organization (*SPARC, COL4A1*) with inflammatory signaling (*IL6*). *FSCN1* also connects into this module through *SPARC*, suggesting a role in cytoskeletal remodeling and cell–matrix interactions during alveolar epithelial transitions. The second cluster involves *MT1E* and *MT1F*, which interact with each other and with *CPVL*, indicating a link between metallothionein-mediated metal ion homeostasis and protease activity. *PRSS23* remains isolated, suggesting a potential independent role in serine protease activity without strong known physical associations to the other proteins in this dataset. Overall, the connectivity pattern points to two functional themes—extracellular matrix/inflammatory regulation and metal ion/protease activity—that may jointly contribute to the cellular processes active across the AT2–transitional AT2–AT1 trajectory.

### Cell fates mediate genetic effects on disease traits

We investigated whether genetic effects on IPF risk are mediated through biased epithelial differentiation decisions at the AT2 transitional point. We considered the following two related trajectories for mediation analysis: AT2–transitional AT2–AT1 and AT2–transitional AT2-transitional AT2-KRT5-/KRT17+. This choice reflects established IPF pathology, in which alveolar type 2 (AT2) cells, normally responsible for regenerating alveolar type 1 epithelium, frequently fail to completely differentiate and instead accumulate in an arrested transitional or aberrant epithelial states. A subset of these cells diverges into a dysplastic lineage marked by KRT17 expression in the absence of KRT5, distinguishing them from classical basal cells. Spatial transcriptomic studies of IPF lungs have shown that transitional AT2 cells exhibit reduced capacity for AT1 differentiation and preferentially populate a non-regenerative KRT5^−^/KRT17^+^ trajectory in diseased tissue [32].

To test whether genetic effects on IPF risk operate in part by biasing epithelial fate decisions, we defined a donor-level quantitative differentiation propensity that captures the net tendency of transitional AT2 cells to progress toward the regenerative AT1 fate versus the aberrant KRT5^−^/KRT17^+^ fate. We estimated both the direct causal effect of gene expression on IPF risk and the indirect effect mediated through differentiation propensity using a linear structural equation model with accumulated genetic effects over a fixed pseudotime interval (Methods). Statistical significance of mediation was assessed using Sobel’s test.

In total, 30 genes exhibited significant mediated effects on idiopathic pulmonary fibrosis (IPF) risk (p < 0.05). Across these genes, greater differentiation propensity toward the AT1 lineage was consistently associated with reduced IPF risk, consistent with the established requirement for completion of AT2→AT1 regeneration after injury [44]. Among these genes, 14 increased differentiation propensity, indicating a shift toward regenerative AT1 commitment rather than the aberrant KRT5^−^/KRT17^+^ fate that accumulates in fibrotic lung niches [45]. Notably, *CLDN3* (Z = 4.37) showed a strong positive mediation effect, consistent with the central role of claudin-family tight junction proteins in alveolar epithelial barrier organization [46]. *MYLK* (Z = 4.83) also promoted AT1 differentiation propensity, consistent with the importance of actomyosin–junctional remodeling during epithelial wound closure and barrier reconstitution [47]. We additionally observed enrichment of lipid-trafficking and surfactant-linked genes among AT1-promoting mediators, including *NPC2* (Z = 2.98). This direction is compatible with the tight coupling between cholesterol/lipid handling, lamellar body biology, and the surfactant metabolic state that must be remodeled during alveolar epithelial regeneration [48].

Two signaling regulators further supported an AT1-biased fate. *VASN* (Z = 3.58), which can antagonize TGF-β signaling by binding and limiting ligand availability, promoted AT1 differentiation propensity [49]. *DUSP6* (Z = 3.14), a negative-feedback phosphatase that dampens ERK pathway activity, also favored the AT1 trajectory, aligning with evidence that ERK signaling operates in tightly regulated feedback loops that can shift between adaptive repair and pathological remodeling depending on magnitude and duration [50]. In contrast, genes biasing differentiation toward the KRT5^−^/KRT17^+^ trajectory were enriched for extracellular matrix (ECM) and mechanotransduction components. *COL4A3* (Z = −3.97) and *COL12A1* (Z = −3.87) were significant drivers of this aberrant fate, consistent with extensive evidence that altered ECM composition and mechanics are not merely end-products of fibrosis but actively reshape cell behavior and differentiation programs [51]. *ANKRD1* (Z = −5.23), a mechanosensitive nuclear factor, emerged as a top driver, providing a direct link between a stiffened niche and transcriptional programs that accompany failed epithelial regeneration [52].

Although endoplasmic reticulum stress is a recognized feature of IPF alveolar epithelium, our mediation results resolve directionally distinct proteostasis programs across fates [53]. The chaperone *DNAJB1* favored AT1 commitment, consistent with a repair-compatible adaptive folding response, whereas *DERL3* (Z = −4.08), a component of the ER-associated degradation machinery, promoted the aberrant KRT5^−^/KRT17^+^ fate, consistent with a shift toward degradation-dominant proteostasis under high stress burden.

Finally, we observed that *FOSL1* promoted AT1 differentiation propensity in our model, despite prior work associating sustained AP-1 activity with fibrotic remodeling and epithelial plasticity programs. Rather than contradicting these studies, this direction is consistent with emerging evidence that AP-1 activity is a core regulator of the injury-induced AT2 activation program and the KRT8^+^ transitional state that precedes AT1 differentiation; in this framework, AP-1 can be required for early activation and chromatin remodeling during regeneration, while persistent or dysregulated AP-1 signaling (for example, in the setting of impaired negative feedback through ERK regulators such as *DUSP6*) may contribute to pathological remodeling [54]. We also nominate *TCIM* and *NCKAP5* as candidate fate regulators with large mediated effects but limited prior characterization in pulmonary fibrosis, highlighting potentially underappreciated genetic control points for epithelial fate bias in the fibrotic lung.

## Discussion

In this study, we highlighted the idea of continuous cell type accounting for cellular variability and stepped further from the existing MR frameworks that consider exposure varying over discrete time. We mapped expression quantitative trait loci (eQTL) effects in the cis-region as functional parameters, which we called “pseudotemporal eQTLs”, showing regulatory effects brought by genetic mutations change continuously along the cellular trajectory. For eQTLs of constant effects across pseudotime we leveraged external bulk-eQTL information to enhance the power. Furthermore, we employed significant pseudotemporal eQTLs as instrumental variables to infer (pseudo-)time varying causal relationships between gene expression and complex traits. To address challenges inherent to scRNA-seq data, such as sparsity and high variability, we incorporate an empirical likelihood-based inference method, which is non-parametric and self-normalized. To rule out the effect brought by a possible mediator that drives the trajectory branchpoint (or the cell fate itself), we implemented summary statistics based MVMR-Egger method with the pointwise effects.

Applying this framework to lung scRNA-seq data and IPF GWAS, we identified causal genes that converge on key processes underlying fibrotic remodeling. Genes with pseudotemporal eQTL effects highlighted the importance of extracellular matrix organization, TGF-β signaling, and immune–epithelial crosstalk, all of which are central to IPF pathology. Moreover, trajectory-specific analyses revealed distinct signatures: immune activation, proteostasis imbalance, and cytoskeletal remodeling in the AT2–AT1 path versus aberrant ciliogenesis and cytoskeletal reprogramming in the AT2–KRT5^−^/KRT17^+^ path. These findings provide evidence that genetic effects influence not only whether transitional AT2 cells differentiate successfully into AT1 cells but also whether they become trapped in maladaptive states that sustain fibrotic progression.

Limitations of this framework are as follows. In our application, the inferred causal relationships are sensitive to both data sparsity and the accuracy of pseudotime estimation, meaning that insufficient cell coverage or invalid trajectory inference can twist the conclusions. Moreover, the pointwise validity of the estimated time-varying causal effects depends on the statistical significance of the eQTLs serving as IVs at each pseudotime point. This requirement can be problematic because cis-eQTLs often share similar pseudotemporal patterns due to linkage disequilibrium (LD), which reduces the independence of IVs and may inflate apparent consistency in causal effects. Consequently, both the robustness of pseudotime inference and the genetic correlation structure need to be carefully considered when interpreting results.

Future work can extend this framework in several ways. First, incorporating multi-omic single-cell data such as scATAC-seq or multiome profiling could allow pseudotemporal eQTLs to be directly linked with chromatin accessibility and enhancer activity, providing a more mechanistic view of regulatory dynamics. Second, applying the framework to larger and more diverse cohorts will improve power to detect weaker or cell-type–restricted effects and enable examination of inter-individual variability in genetic control of trajectories. Finally, extending the model to explicitly account for branching trajectories and trans-eQTLs could capture a wider spectrum of regulatory influences on cell fate. Together, these advances would broaden the utility of pseudotemporal causal inference and help translate dynamic genetic regulation into actionable insights for IPF and other fibrotic diseases.

## Methods and Materials

### Single-cell RNASeq data on human lung

We use the scRNA-seq data from the published study [6], which generated single-cell transcriptomic and genotype data from human lung tissue to investigate cell-type–specific gene regulation and its role in pulmonary fibrosis. The study analyzed lung samples from 66 individuals with fibrotic lung disease and 48 unaffected controls, profiling 38 distinct lung cell types. Using a pseudo-bulk eQTL mapping framework, the authors identified both shared and cell-type–specific eQTLs, and further characterized disease-interaction eQTLs whose regulatory effects differ between fibrotic and healthy states. These interaction eQTLs were found to be highly cell-type specific and enriched in cell types with disease-related transcriptional changes.

We used the top 2000 highly variable genes (HVG) from the Seurat object for epithelial cells from [6] and limit the cell population to the ones we modeled. We extracted the gene–cell matrix and applied two expression filters to reduce sparsity and remove lowly expressed features: (i) we required each retained gene to have zeros in ≤10% of cells (i.e., expressed in ≥90% of cells), computed as the per-gene count of zero entries; and (ii) we required an absolute mean normalized expression ≥0.01 across cells. After the filtering we have 1052 genes and 69463 cells for eQTL mapping. To reconstruct differentiation dynamics, we generated a UMAP embedding of cells from the full HVG integrated expression matrix and applied Slingshot with fixed starting states to infer pseudotime trajectories.

### OneK1K B cells

We used the scRNA-seq data from the [5] of human PBMC. We harmonized B-cell labels by collapsing “B intermediate”/ “B naive” to B_IN and “B memory” to B_Mem, then processed raw counts with Seurat using LogNormalize (scale factor = 10,000), selection of the top 500 variable features by VST, and ScaleData on these features. From the scaled RNA assay we extracted a gene–cell matrix for the target cells and retained genes expressed (zero entries) in ≤10% of cells and with mean absolute scaled expression ≥ 0.01. After filtering we have 404 genes and 125825 cells.

### GWAS summary statistics for IPF

We assembled idiopathic pulmonary fibrosis (IPF) genome-wide association study (GWAS) summary statistics from different large-scale resources, including FinnGen Release 12 [20], the Million Veteran Program (MVP)[21], and the Global Biobank Meta-analysis Initiative (GBMI) [22], comprising a total of 13,760 cases and 2,368,967 controls. All datasets underwent a uniform quality-control and harmonization pipeline. We excluded variants with minor allele frequency (MAF) ≤1%, non-biallelic variants, and variants with missing or non-finite effect estimates, standard errors, or allele frequencies. Variants were mapped to the TOPMed reference panel using genomic position and allele information, with allele orientation harmonized across studies and variants exhibiting discordant allele frequencies removed. We assigned rsIDs using dbSNP build 155 and excluded duplicate or inconsistently annotated variants. To ensure comparability across cohorts, we calculated Z-scores as the ratio of effect size to standard error and derived standardized effect estimates and standard errors using allele frequencies and effective sample sizes. We then performed a meta-analysis using the decoupling framework of [55] to account for overlapping samples across studies. Briefly, we estimated the correlation structure among study-specific association statistics from the degree of sample overlap and incorporated these correlations into the covariance matrix of the effect estimates. We subsequently transformed the covariance matrix into a diagonal form while preserving the original effect estimates, yielding adjusted standard errors that appropriately downweighted non-independent contributions arising from sample overlap. The resulting decoupled summary statistics were then combined using inverse-variance weighting.

### Pseudotemporal eQTL mapping

For a list of genes to map sc-eQTL, the goal is to estimate the eQTL effect function *β* (*t* ). The gene expression is a functional response while the genotype is scalar. This is a functional-on-scalar regression, which is well-contexed in functional data analysis, and could be done by OLS for functional data with B-splines and a smooth penalty. The single-cell level eQTL model includes individual-level covariates, such as sex, age, and ancestry PCs for donors, and cell-specific covariates, such as expression PCs, as fixed effects and accounts for intra-individual cell-to-cell variability using a random effect:

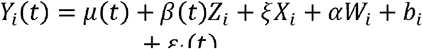

where *Yi*(*t*) denotes the gene expression level at pseudotime *t* from donor *i, Z*_*i*_,*X*_*i*_,*W*_*i*_ denotes the SNP read (0,1,2), individual-level covariates, ancestry PCs respectively.*b*_*i*_ denotes an individual level random effect (intercept) and follow a Normal distribution of *N*(0, *σ*^2^), and *ε*_*i*_ (*t*) denotes a residual error. The functional parameter of interest is *β* (*t*), which we refer as pseudotemporal eQTL. For the interaction pseudotemporal eQTLs, we fit the following model:

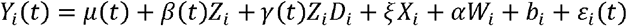

where *D*_*i*_ denotes the donor fibrosis disease status (as control or case).

We have 77 individuals available for trajectory AT2 – Translational AT2 – AT1. After obtaining the pseudotime from Slingshot, the values of pseudotime are rounded by 0.5. We filter out sparse timepoints that there are less than 20 individuals available and exclude genes with fewer than 30 timepoints available. We include cis-SNPs within 1Mb window and exclude “LOWCONF” SNPs. Finally, we apply a 5 cubic B-splines bases with first order difference penalty for the gene expression levels. In practice for the sake of computational efficiency, we firstly regress out the following fixed effects from covariates: gender, age, ethnicity, flow cell ID, processing site, and random effects from the inverse of number of cells per donor and the donor.

We define pseudotemporal eQTL as significant at p-value < 1e-9 level. For eQTLs that are not significant, we fit a constant eQTL effect model as following:

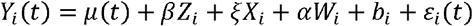

And we utilize an external set of bulk eQTL summary statistics from TOPMed [19], denoted as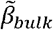, to calibrate 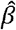 . Although constant eQTLs cannot be used as IVs to estimate the pseudotime-varying causal effect, they still reflect potential mutation effects and increasing power of discovering them is beneficial. After checking for ambiguous SNPs and flipped allele, and harmonizing both sets of summary statistics, we use the method from [41]. Aside from the constant eQTL model, we also fit a bulk model:

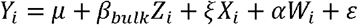

on the single-cell expression data to get 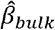. Expression levels are averaged over the pseudotime. We next scale all the estimators as follow:

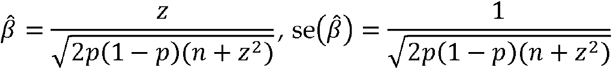

And the calibrated estimator is as follow:

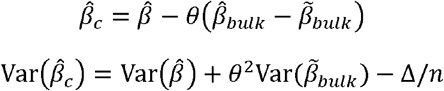

where θ and Δ are functions of residual covariance matrices from the constant eQTL model and the bulk model (details in Supplemental Material).

### Pseudotime varying causal effect estimation

For some gene, denote its estimated pseudotemporal eQTLs as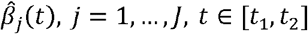 is the pseudotime interval given by Slingshot. Denote the estimated association between SNP and the disease (IPF) as 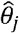 with known se 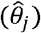. To fulfill the two-sample setting requirement these estimations are from meta-analyzed results from 3 studies [20][21][22]. The main parameter of interest is (1) the (average) pleiotropic effect from the SNP on the disease outcome, denoted as *α*_*0*_, this setup is the same as MR-Egger; (2) the pseudo-temporal causal effect from the gene on the disease outcome, denote as *α* (*t* ). These estimates can be obtained by fitting the following weighted regression, given the core IV assumptions:

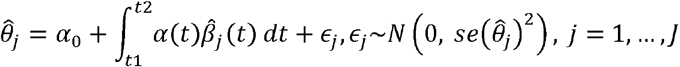

This regression can be fitted using a weighted empirical method [42] [9]. In detail, we can obtain a Karhunen-Loeve representation for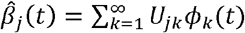, then the regression model can be approximated by

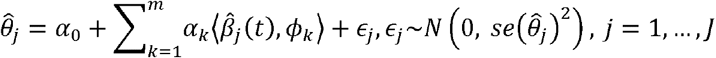

where *αk* = ⟨ *α*(*t*), *ϕ k* ⟩ and *m* is sufficiently large Replace the *ϕ k* with its empirical estimate 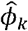, and denote 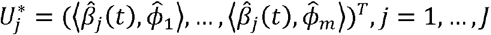. The estimator of*α= α*_*1*_…, *α*_*m*_ depends on *α*_*0*_:

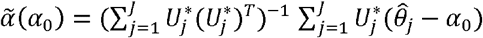

And finally, we estimate *α* (*t*) by

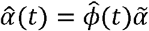

To estimate *α*_*0*_, we use a weighted empirical likelihood method. This is done by maximizing 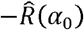, where 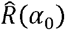 is the following profile empirical likelihood ratio function:

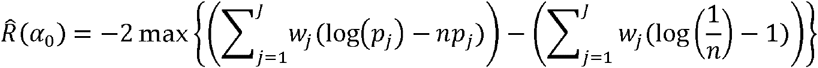

under the constraints

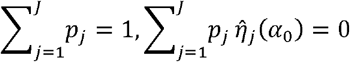

Where 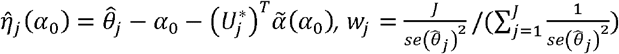 . The confidence interval for 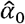 is constructed by 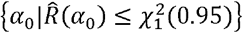.

All the IVs went through LD-pruning with threshold *r*^2^ < 0.1. We considered genes with more than 7 IVs available to ensure identifiability.

### Causal mediation analysis

With the results of inferred two trajectories, AT2–transitional AT2–AT1 and AT2–transitional AT2-transitional AT2-KRT5-/KRT17+, we extract the matrix of cell weights along each lineage with *slingCurveWeights()*, denoted as (W_1*l*_, W_2*l*_), where *l* indexes the cells. We define the propensity difference of each cell as 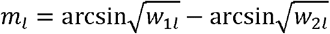 . We also define the weight of each cell as *ω*_*l =*_ *l (m*_*l*_ *> 0*_*1l*_/max (*t*_11_,…, *t*l*L* )+(1-*I*(*m l* >0))*t*_*2*_*l* /max (*t*_21_,…, *t*_2*L*_ to describe the relative position of each cell on the “preferred” trajectory. Finally we define the donor-level cell differentiation propensity as

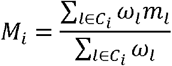

where *C,i* denotes the set of cells belong to donor *i*. Intuitively, if the value of *M*_*i*_ is positive, that means the AT2 cells in donor *i* tend to become AT1 more; otherwise, KRT17^+^/KRT5^−^.

We propose using the following linear structural equation model (LSEM) with accumulated genetic effect for the relationship among functional gene expression, donor-level cell differentiation propensity and IPF status:

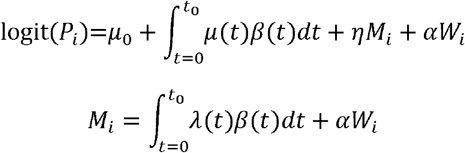

where *P*_*i*_ = P(*Y*_*i*_ = 1) . In our application, *t*_0 =_13.5 which is an approximate critical point of transitional AT2 pre-differentiation status. The mediation effect is defined by the product method, and the standard assumptions of mediation effect can be identified are provided in [43]. This LSEM can be fitted in a common two-step procedure with scalar-on-function regression, and we want to test the hypothesis 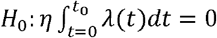 through the Wald-type Sobel’s test:

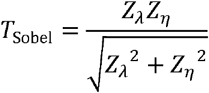

Where 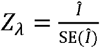 . In detail, 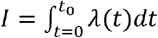, where λ(*t*) was fitted using a 5-dimensional B-spline basis expansion 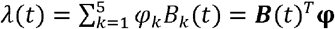 . Hence 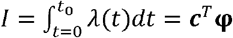, where each element of is computed numerically by trapezoidal quadrature over [0,*t*_0_]. 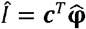 . Note that the Sobel test typically uses the standard normal distribution as the reference distribution to compute the p-value and hence it is conservative for testing the mediation effect.

## Notes

### Competing Interest Statement

The authors have declared no competing interest.

